# Multi-omics investigation of *Clostridioides difficile*-colonized patients reveals protective commensal carbohydrate metabolism

**DOI:** 10.1101/2021.08.24.457492

**Authors:** Skye R. S. Fishbein, John I. Robinson, Tiffany Hink, Kimberly A. Reske, Erin P. Newcomer, Carey-Ann D. Burnham, Jeffrey P. Henderson, Erik R. Dubberke, Gautam Dantas

**Author notes:** <u>Data deposition</u> Metagenomics reads were deposited under BioProject accession number PRJNA748262 and RNA sequencing reads were deposited under BioProject accession number PRJNA748261. <u>Disclaimer</u> The conclusions from this study represent those of the authors and do not represent positions of the funding agencies.

## Abstract

*Clostridioides difficile* infection (CDI) imposes a substantial burden on the health care system in the United States. Understanding the biological basis for the spectrum of *C. difficile*-related disease manifestations is imperative to improving treatment and prevention of CDI. Here, we investigate the correlates of asymptomatic *C. difficile* colonization using a multi-omics approach, comparing the fecal microbiome and metabolome profiles of patients with CDI versus asymptomatically-colonized patients. We find that microbiomes of asymptomatic patients are significantly enriched for species in the class Clostridia relative to those of symptomatic patients. Asymptomatic patient microbiomes were enriched with fucose, rhamnose, and sucrose degradation pathways relative to CDI patient microbiomes. Fecal metabolomics corroborates this result: we identify carbohydrate compounds enriched in asymptomatic patients relative to CDI patients, and correlated with a number of commensal Clostridia. Further, we reveal that across *C. difficile* isolates, the carbohydrates rhamnose and lactulose do not serve as robust growth substrates *in vitro*, corroborating their enriched detection in our metagenomic and metabolite profiling of asymptomatic individuals. We conclude that in asymptomatically-colonized individuals, carbohydrate metabolism by other commensal Clostridia may prevent CDI by inhibiting *C. difficile* proliferation. These insights into *C. difficile* colonization and putative commensal competition suggest novel avenues to develop probiotic or prebiotic therapeutics against CDI.

## Introduction

*Clostridioides difficile* infection (CDI) remains a significant cause of morbidity and mortality in the health care setting and in the community [1]. Antibiotic treatments, among other risk factors associated with weakened colonization resistance, increase susceptibility to CDI [2, 3]. *C. difficile* residence in the human gastrointestinal (GI) tract may result in a spectrum of disease, from asymptomatic colonization to severe and sometimes fatal manifestations of CDI [4]. Diagnosis of CDI relies on detection of the protein toxin, most commonly by enzyme immunoassay (EIA), or the detection of the toxin-encoding genes, by nucleic acid amplification test (NAAT). These diagnostic tools serve as rough benchmarks for assessing severity of disease. Discrepancies between the results of these assays, as in the case of patients with clinically significant diarrhea (CSD) who are EIA negative (EIA-) but NAAT positive for toxigenic *C. difficile* (Cx+), highlight the complexity of states in which *C. difficile* can exist in the GI tract. Clarifying the biological differences between asymptomatic colonization (Cx+/EIA-) and CDI (Cx+/EIA+) will be critical for identifying mechanisms of colonization resistance, and for defining novel probiotic or prebiotic avenues for treatment or prevention of CDI [5, 6].

*C. difficile* enters the GI tract as a spore, germinates in the presence of primary bile acids, and replicates through consumption of amino acids and other microbiota/host-derived nutrients [7]. It is not a coincidence that many of these metabolic cues are characteristic of a perturbed microbiome [8, 9]. The hallmark of *C. difficile* pathogenesis is the expression of the toxin locus encoded on the *tcd* operon; this locus is tightly regulated by nutrient levels [10]. Correspondingly, it is hypothesized that an environment replete of nutrients induces toxinogenesis, allowing *C. difficile* to restructure the gut environment and acquire nutrients through inflammation [11, 12]. Patients who are colonized but have no detectable *C. difficile* toxin in their stool suggests that these patients’ microbiomes may be less permissive towards CDI development. Identification of metabolic traits within the microbiome of asymptomatic, *C. difficile*-colonized patients could reveal a number of potential therapeutic pathways towards precise amelioration of symptomatic *C. difficile* disease.

A multitude of probiotic and prebiotic approaches have demonstrated efficacy to curb *C. difficile* proliferation *in vivo* [6, 13, 14]. While restoration of the microbiota through fecal microbiota transplantation can provide colonization resistance [15], the molecular mechanisms of how this resistance is conferred remain unclear. Recent studies using a murine model of infection have indicated that the administration of carbohydrates (both complex and simple) in the diet can be used to curb or prevent CDI [16-18].

Paradoxically, integrated metabolomics and transcriptomics data collected during murine *C. difficile* colonization indicates that simple carbohydrates are imperative for pathogen replication [11]. It is critical to understand the mechanism by which catabolism of specific carbohydrates could inhibit *C. difficile* proliferation in the human GI tract.

Here, we perform a multi-level investigation of two relevant patient populations, those colonized with *C. difficile* but EIA negative (asymptomatically colonized) and those who are EIA positive (CDI) to understand the microbial and metabolic features that may underlie protection from CDI. First, we use microbiome analyses to identify a number of non-*C. difficile*, clostridial species that are negatively correlated with *C. difficile* in asymptomatically-colonized individuals. Secondly, interrogation of a metabolomics dataset from the same patient population [19] reveals increased abundance of a number of carbohydrate metabolites in asymptomatic patients. Finally, we show that these metabolites enriched in asymptomatically-colonized individuals are largely non-utilizable by *C. difficile* isolates. Together, these datasets reveal that asymptomatically-colonized patients are defined by an interaction of clostridial species and carbohydrate metabolites that may serve as a last-line of resistance against CDI in colonized patients.

## Results

The clinical outcome of *C. difficile* colonization of a host is heavily influenced by taxonomic and metabolic constituents of the microbiome. While colonization in many cases can lead to CDI, evidence of cases of colonization without CDI implies some level of protection from outright disease. We sought to identify biological features that distinguish asymptomatically-colonized patients from those with CDI [20].

In a retrospective human cohort of 102 patients with clinically significant diarrhea (CSD), two groups of patients were identified: those diagnosed with CDI (Cx+/EIA+) or those asymptomatically -colonized (Cx+/EIA-), as defined previously [19]. Because antibiotics are a well-known risk factor for CDI, we analyzed previous antibiotic orders (within one month prior to diagnosis) for patients in the Cx+/EIA-and Cx+/EIA+ cohort, as a proxy for antibiotic exposure (Supplementary Table 1). Fitting antibiotic exposure to a multivariate stepwise logistic regression model (McFadden’s R2=0.306) revealed that CDI was significantly associated with cephalosporin exposure (P=2.70E-04; Supplementary Table 2). Analysis of potential antibiotic exposures in our patient cohort confirms the well-known risk that antibiotics pose to CDI [21, 22].

Antibiotics increase susceptibility to CDI through disruption of colonization resistance, mainly conferred to the host via the gut microbiome [23]. We hypothesized that our asymptomatic patients would have increased microbiome-mediated colonization resistance relative to CDI patients. To determine the microbial correlates of disease state, we performed shotgun metagenomic sequencing on patient stool samples from the asymptomatic (n=54) and CDI (n=48) groups. We examined community structure in stool metagenomes and found that there was no significant difference (Wilcoxon rank-sum, *P*=0.78). in alpha-diversity (Shannon diversity) between patient groups (Figure 1A). In addition, we interrogated beta-diversity (Bray-Curtis dissimilarity) between microbiomes of asymptomatic and CDI patients and found no clustering by EIA status (PERMANOVA, P=0.69) or levels of *C. difficile* (Figure 1B). Previous comparative microbiome studies have revealed phylum-level differences in CDI cases versus controls not colonized with *C. difficile* [24]. In contrast, we found no significant differences in relative abundance of bacterial phyla between asymptomatically-colonized patients and patients with CDI (Supplementary Figure 1B).

**Figure 1.**
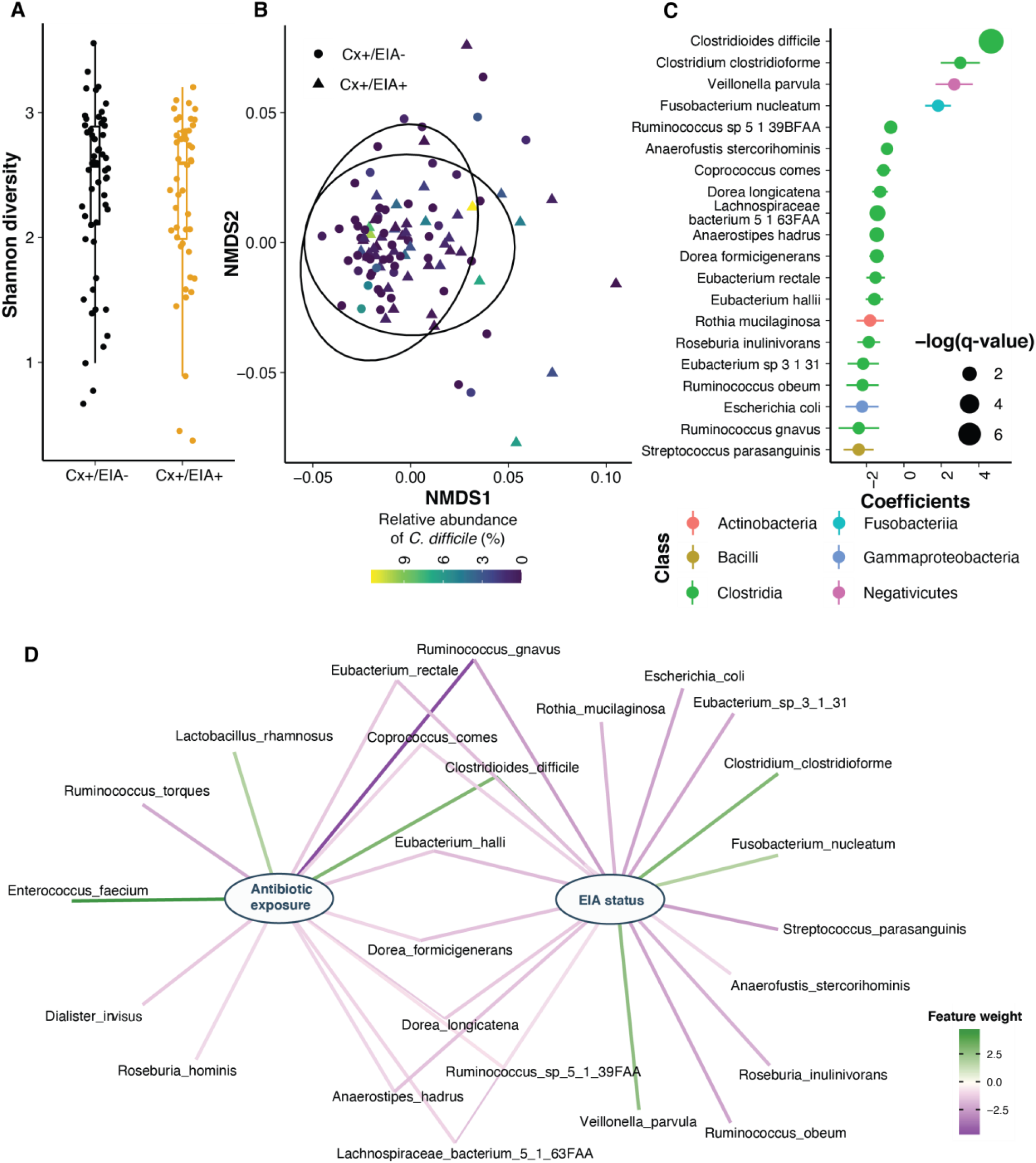
Clostridial taxa enriched in asymptomatic patients. A) Shannon diversity was quantified using stool microbiomes from asymptomatically-colonized (EIA-) and CDI (EIA+) patients. Diversity between groups was not significantly different, as measured by a Wilcoxon rank sum (P=0.78). B) Non-metric multi-dimensional scaling (NMDS) ordination of Bray-Curtis dissimilarity measurements between stool microbiomes. Colors indicate relative abundances of *C. difficile* in each microbiome. Groups were not significantly different as measured by a permutation test of between group dispersion (P=0.69) C) Significant microbial taxa associated with disease state, where a positive coefficient is strongly associated with CDI state. Colors indicate taxonomic Class of microbial feature, and size of circle corresponds to magnitude of statistical significance. Features with q-value of < 0.25 were plotted. D) Network of features associated with antibiotic exposure or EIA status. Species nodes are connected to metadata nodes by edged colored with the feature weight (coefficient) computed from MaAsLin2.

Instead, we hypothesized that differences between these states may manifest at higher resolution. We used a multivariable regression model, as implemented by MaAslin2[25], to identify microbial taxa predictive of either group. *C. difficile* was the strongest predictor of CDI state, whereas non-*C. difficile* Clostridia were predictive of asymptomatic state (Figure 1C, FDR < 0.25; Supplementary Table 3).

Correspondingly, we saw increased *C. difficile* relative abundance in CDI patients and increased levels of a number of non-*C. difficile* clostridial species, including *Eubacterium* spp., *Dorea* spp., and *Lachnospiraceae* spp. in asymptomatic patients (Supplementary Figure 1C). Given that *C. difficile* levels were an overt predictor of CDI, we analyzed patient microbiomes regardless of state to understand microbial features that might correlate with *C. difficile*. Using CoNet [26], a software package that employs multiple measures of correlation to define microbial networks, we found that *C. difficile* anti-correlated with a number of previously identified clostridial taxa (Supplementary Figure 1D). Finally, given the differences in antibiotic exposure in these cohorts, we also interrogated taxonomic features that were predictive of antibiotic exposure. Interestingly, we found that taxonomic features that were predictive of CDI state were also associated with antibiotic exposure (Figure 1D). Our data indicates that patients with *C. difficile* colonization or CDI do not have grossly different gut microbiome community structures but instead have distinctive alterations in a subset species from class Clostridia in the microbiota.

Colonization resistance is often conferred through the presence of commensals that outcompete pathogens in a metabolic niche of the gut [27]. To identify metabolic pathways in other clostridia that might enable them to outcompete *C. difficile*, we defined metabolic potential in patient microbiomes using HUMAnN2 to quantify microbial pathway abundances. In line with our taxonomic analysis, we found no significant differences in alpha- or beta-diversity between overall metabolic pathway composition in the two patient microbiome groups (Supplementary Figure 2A,B). Therefore, we trained an elastic net model to identify specific pathways associated with disease (Figure 2A). We found a number of carbohydrate degradation pathways and amino acid biosynthetic pathways associated with the asymptomatically-colonized (Cx+/EIA-) patients, including ‘sucrose degradation III’, ‘fucose and rhamnose degradation’, and ‘L-methionine biosynthesis I.’ Investigation of the genera that encode such pathways revealed that the sucrose degradation III pathway was increased in asymptomatic patients, largely due to *Blautia* spp. and *Faecalibacterium* spp., among a number of other Firmicutes genera (Figure 2B). Interestingly, the fucose and rhamnose degradation pathways were entirely defined by *Escherichia spp*., presumably *E. coli*. This suggests that metabolic functions such as fucose/rhamnose degradation may be confined to a smaller number of taxa than carbohydrate degradation pathways such as sucrose degradation. Our metabolic pathway analyses highlight differentially abundant carbohydrate degradation processes that could contribute to colonization resistance against *C. difficile* in patient microbiomes.

**Figure 2.**
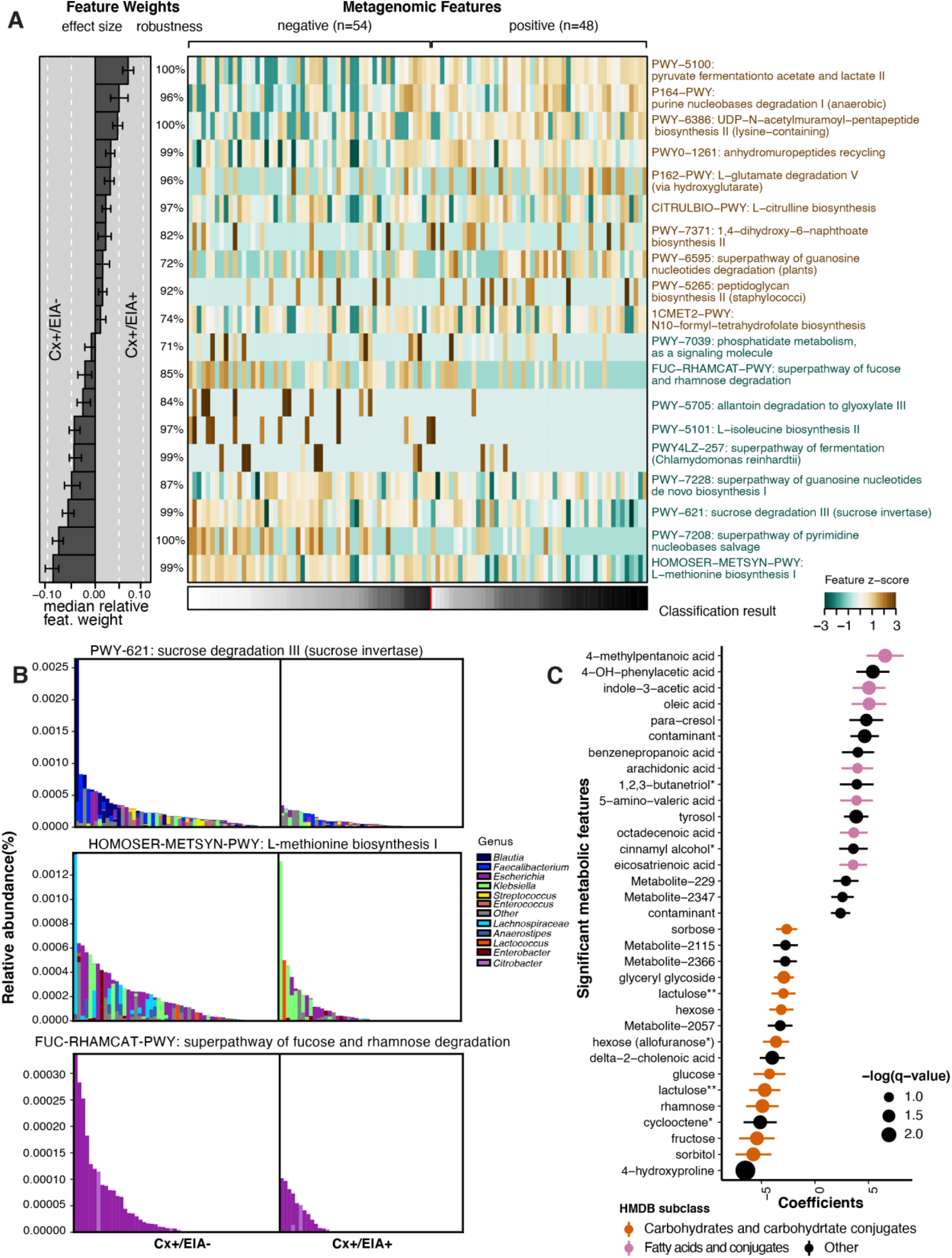
Carbohydrate metabolic processes present in asymptomatically-colonized patient microbiomes. A) Feature weights for elastic net model derived from metabolic pathways abundances. Mean-prediction AUC for the elastic net model was 0.825. B) Relative abundance of taxa in pathways associated with asymptomatic patients. C) Significant metabolites associated with disease state, where a positive coefficient is strongly associated with CDI state. Colors indicate human metabolic database (HMDB) sub-classification, and size of circle corresponds to magnitude of statistical significance. * indicates closest potential match; ** indicates two peaks from the same compound; contaminant indicates mass spectrometry contaminant.

To define metabolic determinants of protection from *C. difficile* in asymptomatic patients, we leveraged metabolomics profiles of these patients’ stools [19]. Ordination of Euclidean distances between Cx+/EIA- and Cx+/EIA+ stool metabolomes revealed no significant differences in metabolome structure (Supplementary Figure 2C). We again used MaAslin2 to determine metabolites associated with each disease state. Consistent with previous analysis, a number of end-product Stickland fermentation metabolites (4-methypentanoic acid and 5-aminovalerate) were associated with CDI patients. While we found that 4-hydroxyproline was the strongest predictor of asymptomatically-colonized patients, many of the significant metabolites that were associated with asymptomatic patients were predicted to be carbohydrates (Figure 2C, FDR < 0.25; Supplementary Table 4). Putative metabolite identities were initially annotated by matching metabolite spectra to the NIST14 GC-MS spectral library. The preponderance of carbohydrates in asymptomatic patients and the substantial similarity of carbohydrate spectra prompted us to rigorously validate the identities of these metabolites by comparing EI spectra and GC retention times against authentic standards, where commercially available (Supplementary Table 2, Supplementary Figure 3). These data reveal a carbohydrate signature that is depleted in CDI patients.

Notably, fructose and rhamnose are either substrates or products of the ‘sucrose degradation III’ and ‘fucose and rhamnose degradation’ pathways, which we found to be enriched in asymptomatic patients. The co-occurrence of these microbial pathways and their corresponding metabolites suggest that the presence of a commensal carbohydrate metabolism that could antagonize *C. difficile* pathogenesis.

We hypothesized that the differential abundance of identified stool metabolites in these patient cohorts is related to the metabolism of specific microbes in their microbiomes. We performed a sparse partial least-squares-discriminatory analysis (sPLS-DA) with the mixOmics package to define relationships between the most predictive features of patient metabolomes and microbiomes. Our final model (Figure 3) contained one latent component, composed of 15 metabolites and 20 microbial species. Of the strongest metagenomic variable weights, four out of five species (*C. difficile*, a *Lachnospiraceae spp*., *Anaerostipes hadrus*, and *Clostridium clostridioforme*) were also significantly associated with an EIA state (Figure 1). In the metabolomics block of the latent component (Figure 3A), the eight highest-weighted metabolites were also discovered by previous analyses (Figure 2). Using the variables defining the latent component, we performed correlational analyses (Figure 3B) and found a number of striking correlations. *C. difficile* and Stickland metabolites (5-amino-valeric acid and 4-methylpentanoic acid)[19] were positively correlated, whereas *C. difficile* had negative relationships with fructose, rhamnose, and hydroxyproline. Moreover, we found that carbohydrates, such as fructose and rhamnose, were correlated with the presence of commensal *Lachnospiraceae* and *Streptococcus* species; the *Lachnospiraceae* spp. was also previously anti-correlated with *C. difficile* (Figure 1). This network revealed both expected relationships, highlighting the known pathophysiology of CDI, and novel commensal-carbohydrate relationships that define asymptomatic colonization.

**Figure 3.**
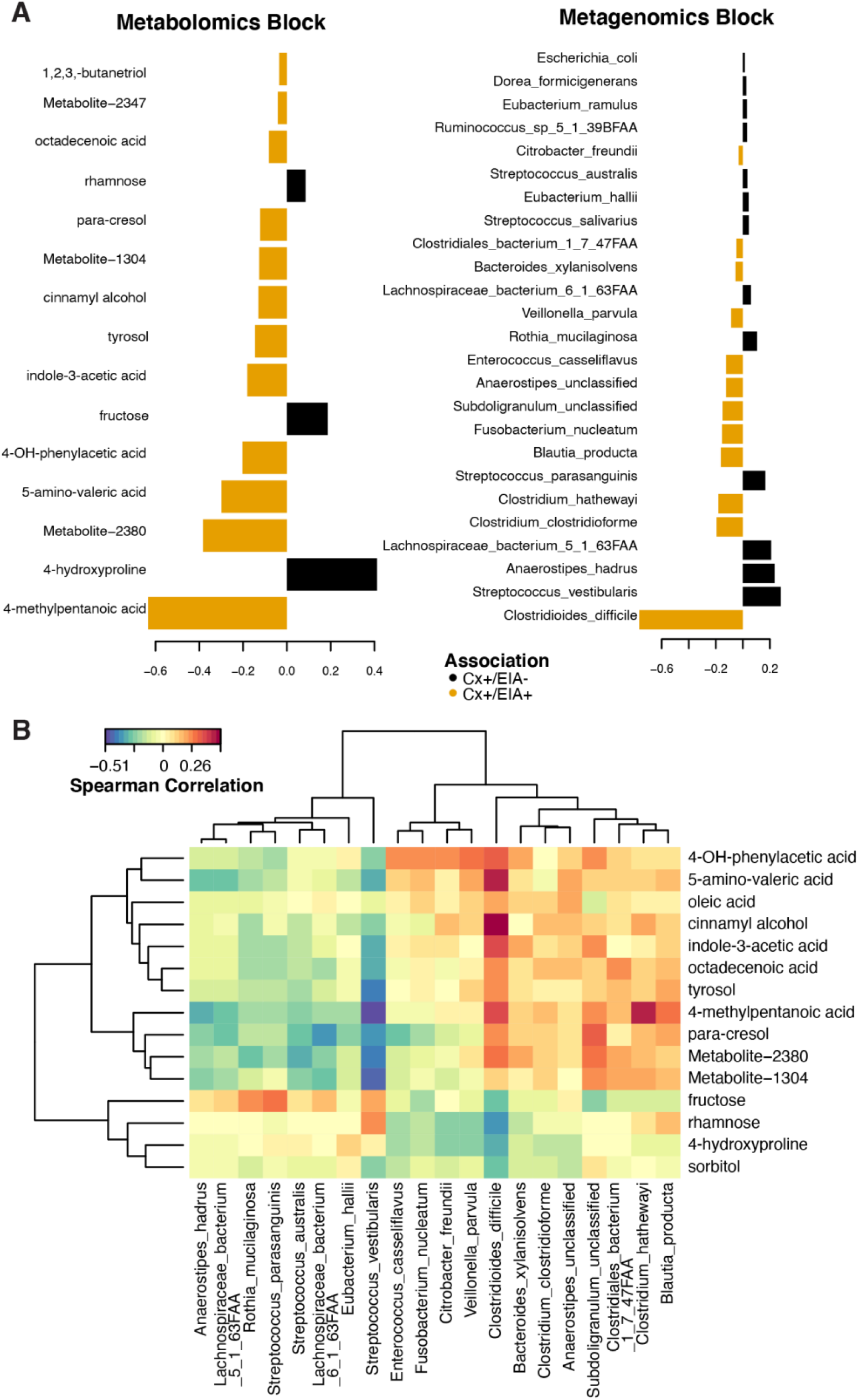
Multi-omics signature of asymptomatic patients. A) Contribution of each variable (microbe or metabolite) to latent component of sparse partial least-squares-discriminatory analysis (sPLS-DA) B) Heatmap of Spearman correlations between metagenomic and metabolomic variables from the latent component.

Our examination of taxa, metabolic pathways, and metabolites revealed that presence of non-*C. difficile* clostridial taxa that might provide colonization resistance against *C. difficile* through their metabolism. We hypothesized that the observed inverse relationship of certain carbohydrates to *C. difficile* might indicate that these metabolites are not be digestible by *C. difficile* or that these metabolites are end-products of a more complex commensal metabolism that is exclusionary to *C. difficile*. Using a set of clinical *C. difficile* isolates cultured from this patient cohort (6 different ribotypes), we examined growth of *C. difficile* on carbohydrates associated with asymptomatic patients. Using a defined minimal media (CDMM[28]), we found that *C. difficile* isolates grew robustly on fructose as expected (median maximum A_600_ of 0.72), but did not proliferate on rhamnose or lactulose (median maximum A_600_ of 0.18 and 0.20 respectively). Notably, in the case of sorbitol, we found that a subset of strains, including the reference strain VPI1064, grew to a maximum A_600_ of greater than 0.35 (Figure 4A,B). Given that we had found sucrose degradation as a metabolic pathway enriched in asymptomatic patients, we hypothesized that *C. difficile* would be unable to use this carbohydrate. Indeed, when grown on sucrose as the sole carbon source, strains achieved a median maximum A_600_ of ∼4.2-fold less than that of growth on fructose.

**Figure 4.**
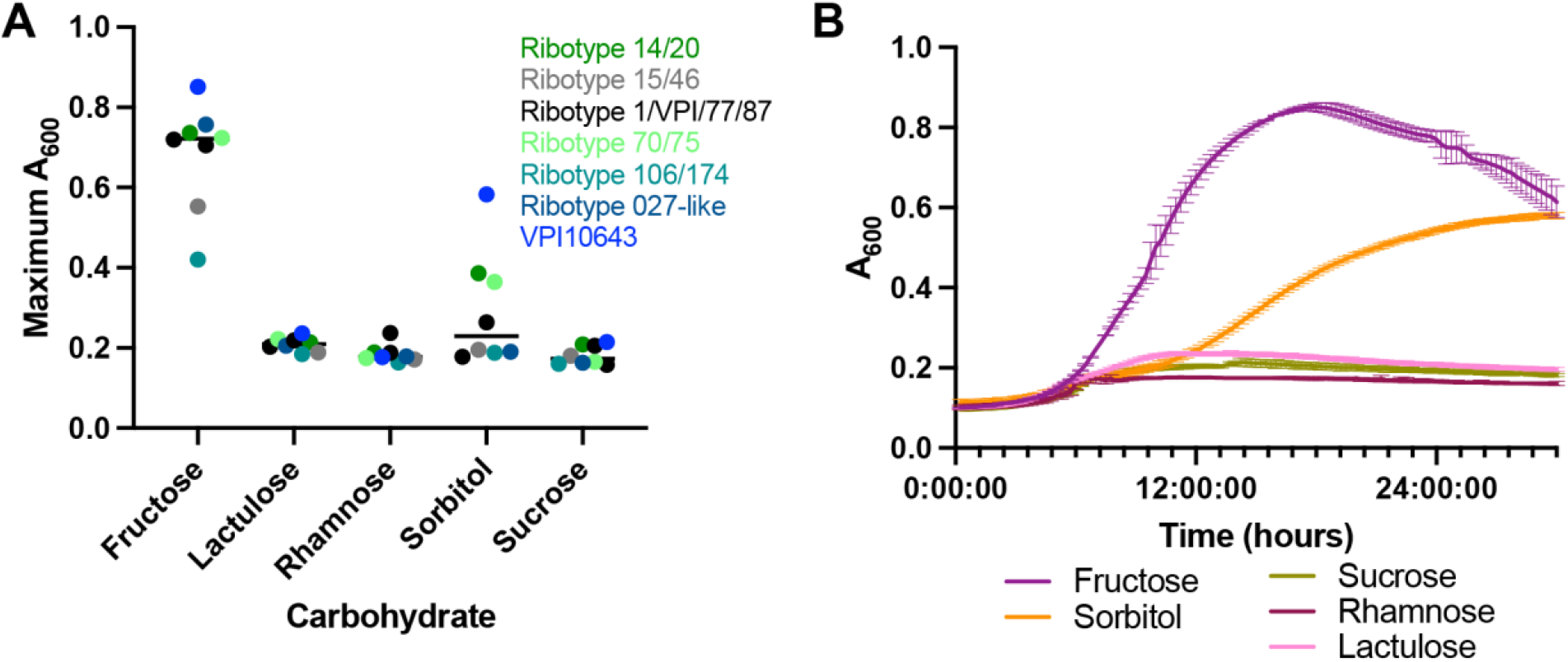
Influence of carbohydrates on growth. A) Clinical isolatesand reference strain grown in *C. difficile* minimal medium (CDMM) with equimolar amounts of each carbohydrate added. Growth was measured by taking maximum absorbance values over 24 hours. Each point represents the mean of two technical replicates of a unique isolate. B) Growth of *C. difficile* VPI 10463, from same conditions in (A). Data in (A-B) are representative of multiple experiments.

Though *C. difficile* cannot grow on rhamnose as the sole carbohydrate, in other organisms rhamnose has substantial transcriptional influence over carbon catabolite gene clusters [29, 30]. We wanted to rule out the possibility that rhamnose may impact *C. difficile* through possibly cryptic transcriptional reprogramming, perhaps contributing to *C. difficile* repression *in vivo*. Accordingly, we performed whole transcriptome RNA sequencing on *C. difficile* cultures exposed to a metabolizable substrate, fructose, or a non-metabolizable substrate, rhamnose. In the presence of fructose, we found 555 genes significantly altered (adjusted p-value <0.05 and |fold-change| > 2)(Supplementary Table 5). Some of the most altered genes were indicative of carbon catabolite repression of sugar transport and upregulation of glycolytic processes to metabolize fructose. In contrast, we found only 3 genes significantly increased in the rhamnose condition (Supplementary Figure 4). The lack of striking systems-level or targeted (toxin expression, sporulation) regulation by rhamnose, and *C. difficile*’s inability to utilize it, leads us to conclude that its association with asymptomatic patients’ microbiomes is not through direct interaction or suppression of *C. difficile*. Instead, we speculate that rhamnose may be the byproduct of a complex commensal metabolism (orchestrated by the non-*C. difficile* clostridia) of other dietary polysaccharide substrates, which could exclude *C. difficile* from the GI tract.

## Discussion

### Antibiotic exposure and loss of clostridial niche related to CDI

Antibiotic treatment is the most well-understood risk factor for CDI [31, 32], and antibiotic exposure in our cohort likely results in loss of the species we find depleted from CDI patients. Here, we confirm that exposure to a number of antibiotics is associated with CDI patients, including cephalosporins (significantly associated) and intravenous vancomycin (weakly associated) patients. Clindamycin and quinolones, two antibiotics also associated with CDI in other human cohorts [33] are likely not significantly associated in our population due to the low prevalence of their exposure. Our microbiome data reveals decreased levels of *Streptococcus, Ruminococcus, Eubacterium*, and *Roseburia* spp. in CDI patients. Findings from both human cohorts and mouse models of antibiotic treatment indicate that a number of these clostridial taxa, also considered major butyrate-producing species, are depleted upon administration of a variety of antibiotic treatments [34, 35]. It is also posited that some of these taxa are integral to protection from CDI [36]. Given the attempts to use FMTs or Firmicutes-enriched probiotics to prevent CDI, we hypothesize that the restoration of lost species from class Clostridia after high-risk antibiotic treatment could be a novel avenue for CDI prevention [37].

The results from our cross-sectional multi-omics profiling of these CDI-related human cohorts could reflect a number of biological scenarios. The observed increased abundance of monosaccharides, in both asymptomatic patients and diarrheal control patients [38], could imply the presence of a more diverse gut microbiota metabolism, wherein commensal microbiota are degrading complex polysaccharides into simpler substrates. While the community-level diversity between the two groups was not strikingly different, the correlation between clostridial taxa and certain carbohydrates further hints at the possibility of the presence of a *C. difficile*-antagonizing metabolism within the microbiome. Finally, we find that many of these identified carbohydrates are not nutrient substrates for a diverse set of clinical *C. difficile* strains, which contrasts with reports of model *C. difficile* strains harboring more flexibility in their ability to acquire and utilize nutrients [23, 39, 40].

On the other hand, identification of the metabolites sorbitol and 4-hydroxyproline in the stool of asymptomatic patients colonized with toxigenic *C. difficile* may indicate low-level inflammation. Recent data indicate that hydroxyproline and sorbitol are host-derived metabolites, reflective of some amount of collagen degradation and toxin-mediated inflammation [12, 41]. In a mouse model of CDI, the presence of sorbitol and mannitol before pathogenesis was interpreted as a “pre-colonized state” [11, 23], and these simple carbohydrates declined as CDI progressed. To restrict our cohort to patients who were not on their way to developing CDI, patients with EIA-stool were excluded if they were subsequently diagnosed with CDI or received empiric CDI treatment within 10 days of initial stool collection [20]. Thus, we conclude that the multi-omics data presented here reflects a more intermediate state where CDI development is hampered by the microbiome or other host-associated factors.

### A combination of commensal-derived and other dietary carbohydrates define asymptomatic patients

Lactulose is a disaccharide product of heat treatment of lactose (a common sugar in dairy products), and also a component of some laxatives [42]}. Yet, patients were screened and excluded from this cohort if they were prescribed laxatives in the 24 hours prior to sample collection. In addition to this screening/exclusion criteria, lactulose is almost exclusively prescribed to liver failure patients, thus it is more likely to be present from consumption of heated milk (containing lactose). Interestingly, lactulose has been associated with a decrease in *C. difficile*-related diarrhea [43] and decreased risk of CDI [43, 44]. Additionally, *in vitro* work demonstrates that addition of “non-digestible” oligosaccharides, such as lactulose, provide a competitive advantage to *Bifidobacterium* spp. over *C. difficile* [5, 45]. Our survey of multiple *C. difficile* ribotypes shows that *C. difficile* cannot utilize lactulose, and agrees with the above *in vitro* competition data. Taken together, these data emphasize the potential for synthetic or natural prebiotic interventions to shift a vulnerable microbiota away from CDI. While we do not recommend lactulose, a laxative, as such a prebiotic, there are a number of other ‘non-digestible’ oligosaccharides that might serve similar purposes in future interventions [45].

We also found an increase in rhamnose in asymptomatic patients, and confirmed that across a number of clinical *C. difficile* isolates, growth with rhamnose as the sole carbohydrate source is severely decreased relative to the robust growth observed with fructose. Transcriptional profiling of rhamnose-exposed *C. difficile* clarifies that this carbohydrate does not induce major systems-level changes to bacterial expression networks or physiology. Rhamnose is a major component of plant and some bacterial cell-wall polysaccharides. The presence of this deoxy sugar could be related to degradation of polysaccharides from either eukaryotic or prokaryotic kingdoms of life [46]. Finally, in the patients asymptomatically-colonized with toxigenic *C. difficile*, metabolic pathway profiling revealed an enrichment of fucose and rhamnose degradation pathways, represented by Enterobacterales taxa. Therefore, we propose that the detected rhamnose is a byproduct of commensal catabolism of more complex polysaccharides containing rhamnose [47, 48]. However, we acknowledge that these data do not necessarily imply that *C. difficile* cannot find usable carbohydrate sources *in vivo* (indeed, glucose and fructose are concomitantly abundant with rhamnose in our data). Future experiments are required to resolve the underlying pathophysiology of the rhamnose-asymptomatic microbiome association.

### Strategies to ameliorate toxigenic *C. difficile* proliferation

Our multi-omics analyses of a colonized asymptomatic patient population support a growing body of literature concerning commensal metabolism as a tool against *C. difficile*. Evidence from both mouse models of disease and human studies indicate that administration of polysaccharides or ‘microbial accessible carbohydrates’ may prevent *C. difficile* proliferation or decrease the risk of CDI [16-18, 43, 49]. Recently, a probiotics-based attempt to design a consortium of mucosal sugar utilizers revealed its ability to decrease *C. difficile* colonization *in vivo* [14], indicating that increasing mucosal metabolism, or carbohydrate catabolism, may be another route to strengthening commensal resistance to *C. difficile*. Given the plethora of prebiotics and probiotics explored in the *C. difficile* field, we emphasize the need for an approach that harnesses both probiotic- and prebiotic-based components to inhibit the proliferation of *C. difficile* and toxin-mediated pathogenesis.

## Acknowledgements

The authors are grateful for members of the Dantas lab for their helpful feedback on the data analysis and preparation of the manuscript. The authors would also like to thank the Edison Family Center for Genome Sciences and Systems Biology staff, Eric Martin, Brian Koebbe, MariaLynn Crosby, and Jessica Hoisington-López for their expertise and support in in sequencing/data analysis.

## Materials and Methods

### Patient cohort analysis

A previous retrospective cohort study was conducted to understand *C. difficile* colonization [20]. For the purposes of this study, data concerning inpatient antibiotic orders were retrospectively collected from the electronic medical informatics database for patients with toxin EIA positive (Cx+/EIA+) stool (n=62) or toxin EIA negative (Cx+/EIA-) stool. Further, the presence of antibiotic orders was classified into three dichotomous groups by timing of exposure: antibiotics in 0-7 days before stool collection (1 week), antibiotics in >7-14 days before stool collection (2 weeks), and antibiotics in >14-30 days before stool collection (1 month). Antibiotics were classified into groups where appropriate. The dichotomous antibiotic usage data was imported into R and analyzed using MaAslin2 with fixed effects set to EIA status, min_abundance=0, min_prevalence=0, normalization=“NONE”, transform=“NONE”, and all other default settings.

### Metagenomics sequencing and analysis of patient stool

Metagenomic DNA was extracted from patient stools as previously described [50]. *C. difficile* was isolated from patient stools as previously described [50]. Illumina libraries of patient stool metagenomic DNA were prepared and pooled as previous described [50, 51]. Fecal metagenomic libraries were submitted as 2×150bp paired-end sequencing on an Illumina NextSeq High-Output platform. Reads were binned by index sequences and reads were trimmed and quality filtered using Trimmomatic v.0.38 [52] to remove adapter sequences and DeconSeq[53] to remove human sequences. Samples that were less than 15% bacterial DNA during initial sequencing were discarded, and all samples were sequenced to a depth of at least 5 million reads.

We performed taxonomic profiling of metagenomic sequences using MetaPhlAn2 [54] and functional pathway profiling using HUMAnN2[55]. Metacyc pathway abundances were normalized to relative abundances using the humann2_renorm.py function. The humann_barplot function was used to assess taxonomic composition of metabolic pathways. Custom python scripts were used to parse metaphlan “_profiled_metagenome.txt” and humann2 “pathwayabundance.txt” files. Data were imported to R to analyze community composition and differential associations. MaAslin2 was used to fit a logistic regression model for microbial taxa/metabolites associated with EIA status as the fixed variable in the 102 patient microbiomes. No normalizations were applied, and data were log-transformed, with all other default settings applied. To analyze HUMAnN2 pathways enriched in cohorts, we used statistical inference of associations between microbial communities and host phenotypes (SIAMCAT)[56], from the siamcat package in R, to fit an elastic net model to the data. We used the following parameters: log.std normalization, 10 folds and 10 resamples for data splitting. The model.interpretation.plot function was used to display features weights for features used in >70% of models generated in training.

### Data analysis

For both microbiome and metabolomics data, the nearZeroVar function of the caret package was used to remove low-prevalent or invariant taxa/pathways/metabolites. These filtered data sets were analyzed for community composition and differential association. Alpha-diversity and beta-diversity were calculated using the vegan package. Bray-Curtis dissimilarity was used as a beta-diversity metric for microbial taxa and metabolic pathways, while Euclidean distance was used as a beta-diversity metric for metabolomes. The MaAslin2 package was used to fit a logistic regression model for microbial taxa associated with EIA status as the fixed variable. No normalizations were applied, and data was log-transformed, with all other default settings applied. This package was also used to fit a logistic regression model for microbial taxa associated with antibiotic exposure. Antibiotic exposure data was converted to a binary variable for this analysis, and microbiome data was treated as above. Finally, MaAslin2 was used to fit a logistic regression model for metabolites associated with EIA status, with the same settings as above.

### Network analysis of microbiome

Filtered relative abundances were loaded into CoNet[26], a program used to determine biological networks. The following options were selected: in the ‘Preprocessing and filter’ menu, ‘col_norm’ was applied; in ‘Methods’ menu, ‘Spearman’, ‘Bray Curtis’ and ‘Kullback-Leibler Divergence’ were applied; in ‘Threshold’, ‘top’ and ‘bottom’ were selected with a threshold of 0.025, ‘force intersection’ was selected; in ‘Merge’, ‘multigraph’ was checked, score merging was performed using ‘mean’ option, and network merge ‘intersection’ option was selected; in ‘Randomization’, 100 iterations was performed, ‘edgeScores’, resampling is ‘shuffle rows’, p-values were merged b ‘geometric mean’, renormalize was checked and pvalue option was ‘benjaminihochberg’ at 0.05. For clarification of relevant interactions, a subnetwork was created where *C. difficile* and first neighbor nodes were selected to visualize the local network.

### Determination of candidate metabolites

Putative identification of metabolites of interest (Supplementary Table 2) was initially performed through spectral matching against the NIST14 electron ionization spectrum library. Several features were previously identified by our group (see Robinson *et al*. 2019). Features predicted to be sugars or sugar alcohols were compared to a panel of authentic standards (D-sorbitol, D-mannitol, D-fructose, L-rhamnose, L-fucose, lactulose, glucose, mannose, D-galactose, D-talose, *myo*-inositol, and L-sorbose). Because isomeric sugars generate very similar spectra, we utilized both spectral similarity and retention time to identify sugar metabolites (Supplementary Figure 2B).

### Multi-omics analysis

The metagenomics relative data was imputed with min(abundance>0)/2, and the metabolomics data was imputed with a value of 1. For both filtered datasets, a centered log-ratio transformation was used to analyze filtered metagenomics and metabolomics datasets above. The mixOmics package in R was used for multi-omics analysis. To avoid over-fitting on the large number of variables in our datasets, we utilized sparse PLS-DA. Briefly, to determine the number of variables from each dataset to keep in the final model, we estimated model error rates for all combinations of seq(5,50,5) variables for both metagenomic and metabolomic datasets, using the function tune.block.splsda (10-fold cross-validation, repeated 50 times, “max.dist” distance metric). We chose to minimize the error with the maximum number of variables, therefore keeping 15 metagenomic variables and 15 metabolomic variables. Spearman correlations were calculated between CLR-transformed microbial taxa and metabolite abundances, from the variables defining the first latent component, and plotted using the cim package.

### Bacteriology and *in vitro* growth assays

*C. difficile* strains were isolated from patient stools by plating on cycloserine-cefoxitin fructose agar as previously described; strains were stored at –80 degrees C [50]. For *in vitro* growth assays, CDMM was prepared as previously described [28] and 20 mM of specified carbohydrates were added. Clinical isolates were inoculated into TY and grown for 16hrs, then washed with PBS and diluted 1:100 into media with different carbohydrates sources. Growth was measured in a shaking, 96-well plate at 37°C for 36 hours.

### RNA sequencing and data analysis

Five mL of each strain (in biological triplicate) were grown to log-phase (OD_600_ ∼ 0.4)in TY and exposed to TY-rhamnose or TY-fructose (with each carbohydrate at 30 mM). Cells were harvested by adding one volume of 1:1(v/v) acetone/ethanol to the culture to arrest growth and RNA degradation.

Sample were spun at 4000 x g for 5 minutes. The cell pellet was washed with 500 µl TE buffer (0.5 M EDTA, 1 M Tris pH 7.4) and spun down to remove the supernatant. The cell pellet was resuspend in one mL Trizol and two rounds of bead-beating at 4500rpm for 45s were performed. 300µl of chloroform was added to the suspension, lysates were vortexed, and centrifuged at 4000 rpm for 10 min at 4°C. The aqueous layer was removed and RNA was precipitated using isopropanol, washed with 70% ethanol, and resolubilized in TE buffer. Total RNA was treated with Turbo DNase (for two rounds of digestion). rRNA depletion was performed using the QiaFastSelect kit (Hilden, Germany), following manufacturer’s instructions. prepared for sequencing using KAPA Stranded RNA-Seq Library Preparation Kit (Millipore Sigma, St. Louis, MO). Libraries were prepared using the rRNA-depleted RNA as input for NEBNext Ultra II RNA Library Prep Kit (NEB, Ipswich, MA). Libraries were pooled and submitted for 2 × 150 bp paired-end sequencing on an Illumina NextSeq High-Output platform at the Center for Genome Sciences and Systems Biology at Washington University in St. Louis.

Raw reads were trimmed using Trimmomatic v. 0.38, and aligned to a *C. difficile* VPI10643 reference genome (GCF_000155025.1) using Bowtie2. SAM files were converted to BAM format and indexed using samtools. Read counts for each gene feature were obtained using the featureCounts function of subread-1.6.5 package. Counts were manually imported into R, and DEseq2 was used to identify differentially expressed gene products in the case of TY-fructose relative to TY and TY-rhamnose relative to TY.

**Supplementary Table 1:** Association of antibiotic orders with disease state. Timing of exposure indicates the time frame before EIA diagnosis. Times of exposure were binned into three dichotomous features (1 week, 2 weeks or 30 days)

| Antibiotic | Coefficient | Standard Error | P-value |
| --- | --- | --- | --- |
| Cephalosporin | 2.68 | 0.74 | 2.70E-04 |
| Fluoroquinolone | 0.19 | 1.09 | 0.86 |
| Carpapenem | 0.34 | 1.12 | 0.76 |
| Metronidazole | 1.11 | 0.95 | 0.24 |
| Vancomycin (intravenous) | 1.44 | 0.95 | 0.13 |
| *Hosmer and Lemeshow Goodness of fit test p=0.0080 |  |  |  |

**Supplementary Figure 1.**
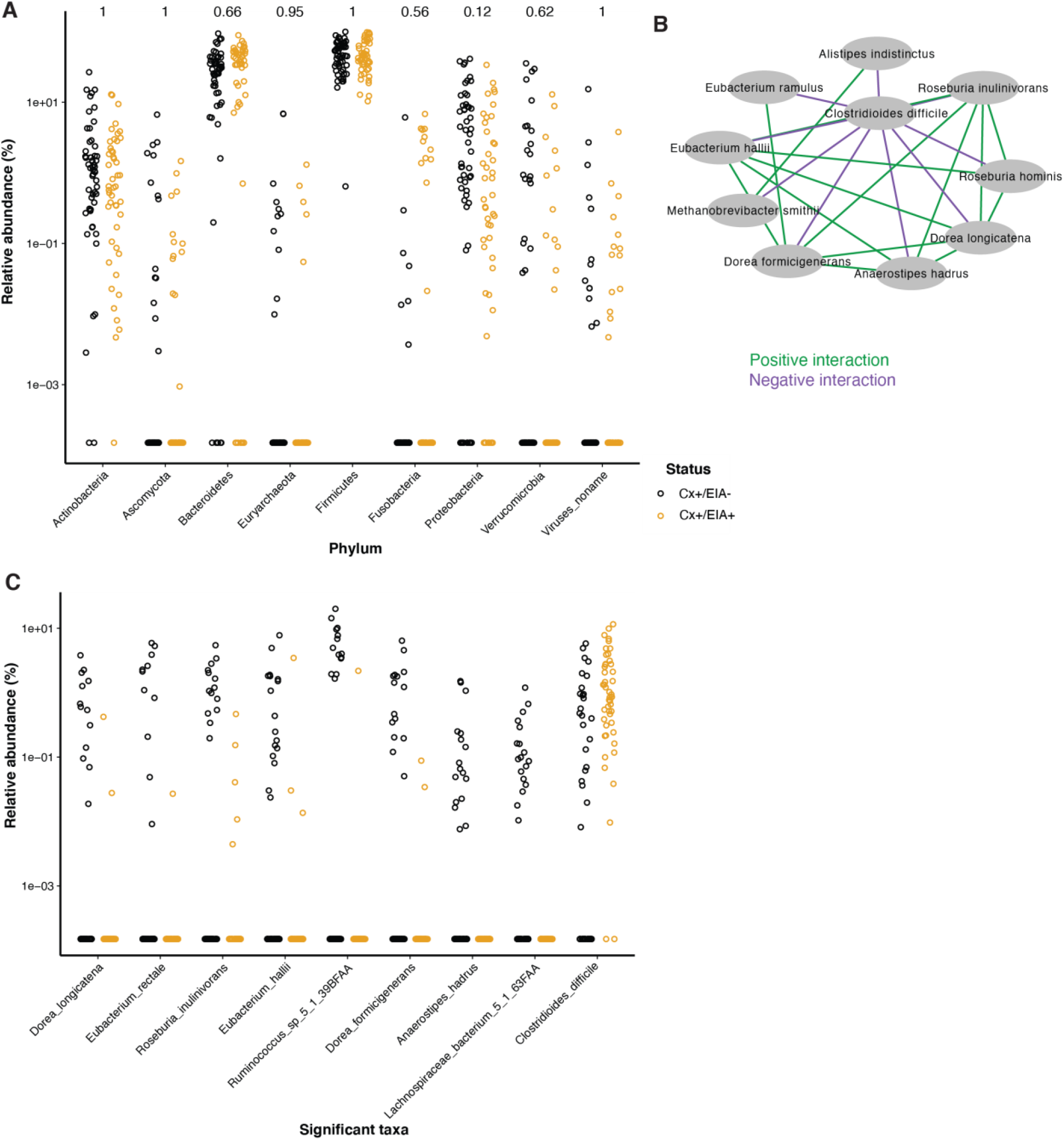
Microbiome configuration of C. difficile-colonized patients. A) Phylum-level comparisons between CDI and asymptomatic patients. P-values above comparisons calculated using a Wilcoxon rank sum test and corrected for multiple testing. B) Network of interactions with *C. difficile*, generated by CoNet, where edges colored in green indicate a positive interaction and edges colored in purple indicate a negative interaction. C) Relative abundance of significantly altered clostridial taxa from MaAsLin2 analysis.

**Supplementary Figure 2.**
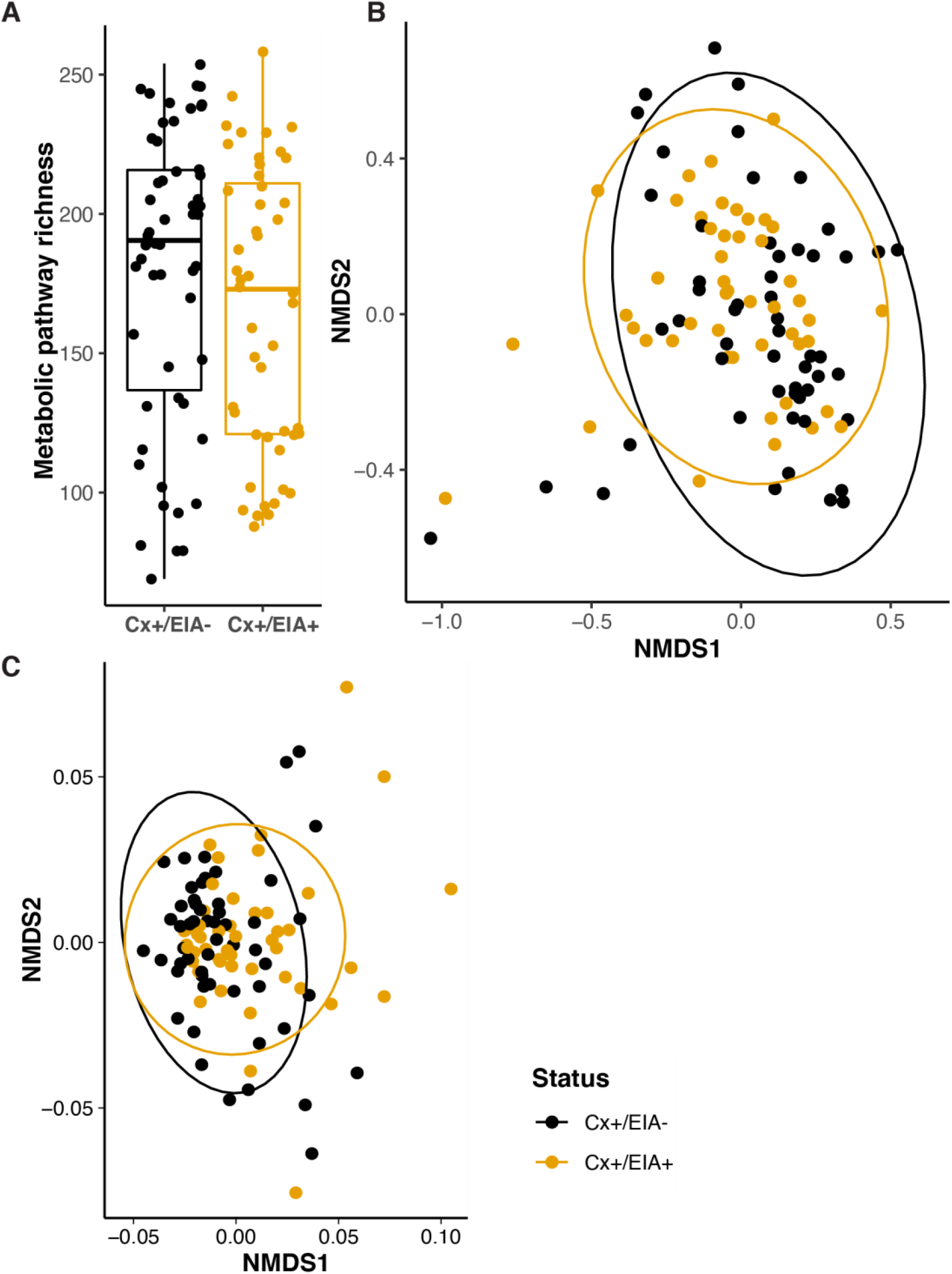
Compositional measurements of metabolic pathways and metabolites. A) Alpha-diversity (measured by richness) of metabolic pathways in patient groups. B) Non-metric multi-dimensional scaling (NMDS) ordination of Bray-Curtis dissimilarity measurements between stool microbiome metabolic pathways. C) NMDS ordination of Euclidean distances between stool metabolomes. A permutation test was used to quantify centroid distances between groups (P=0.01).

**Supplementary Figure 3.**
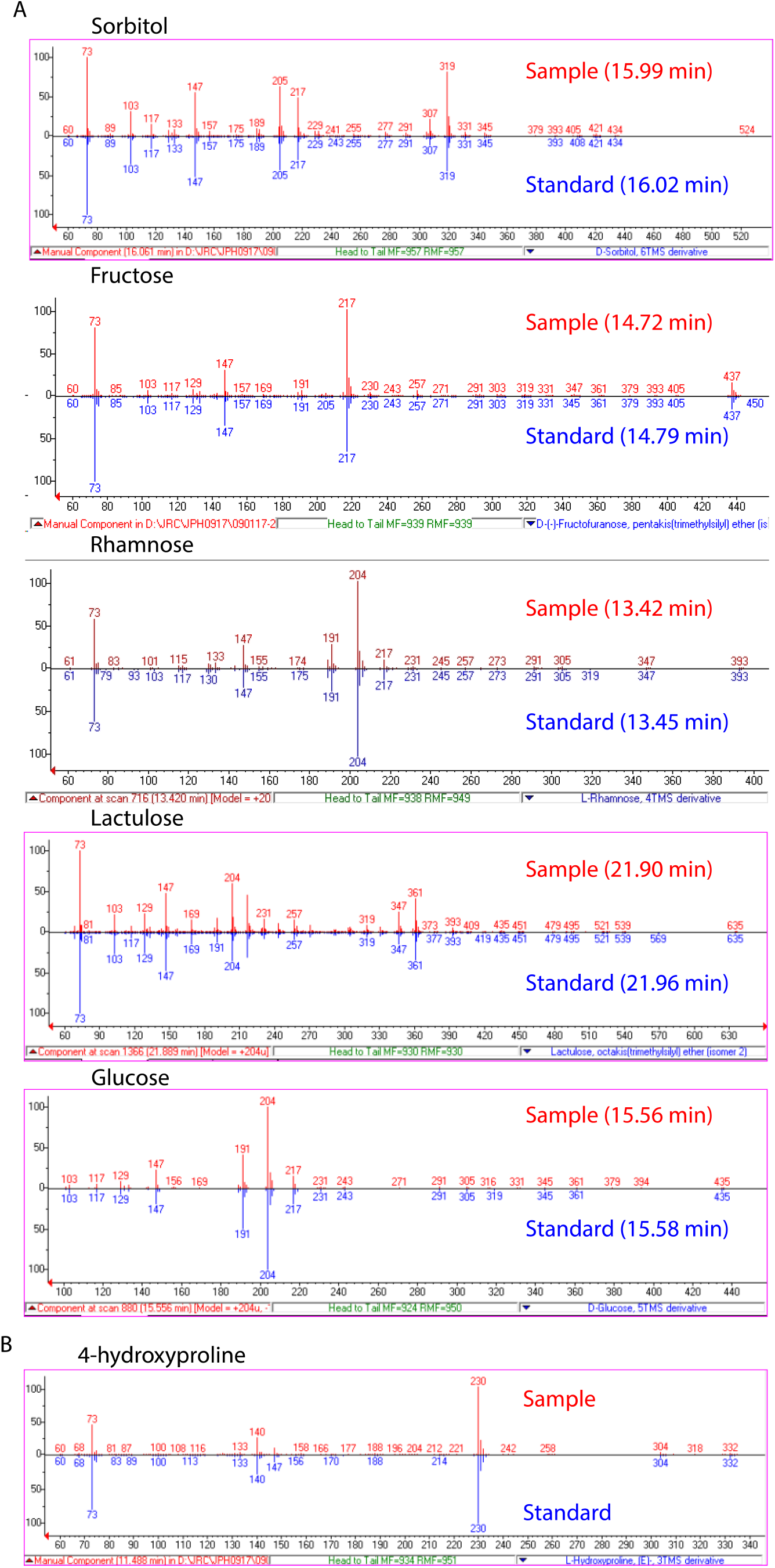
Validation of carbohydrates. A) EI spectra and GC retention times of identified sugar and sugar alcohol features (red) compared to spectra of authentic standards (blue). B) EI spectrum of 4-hydroxyproline feature compared to NIST14 library spectrum.

**Supplementary Figure 4.**
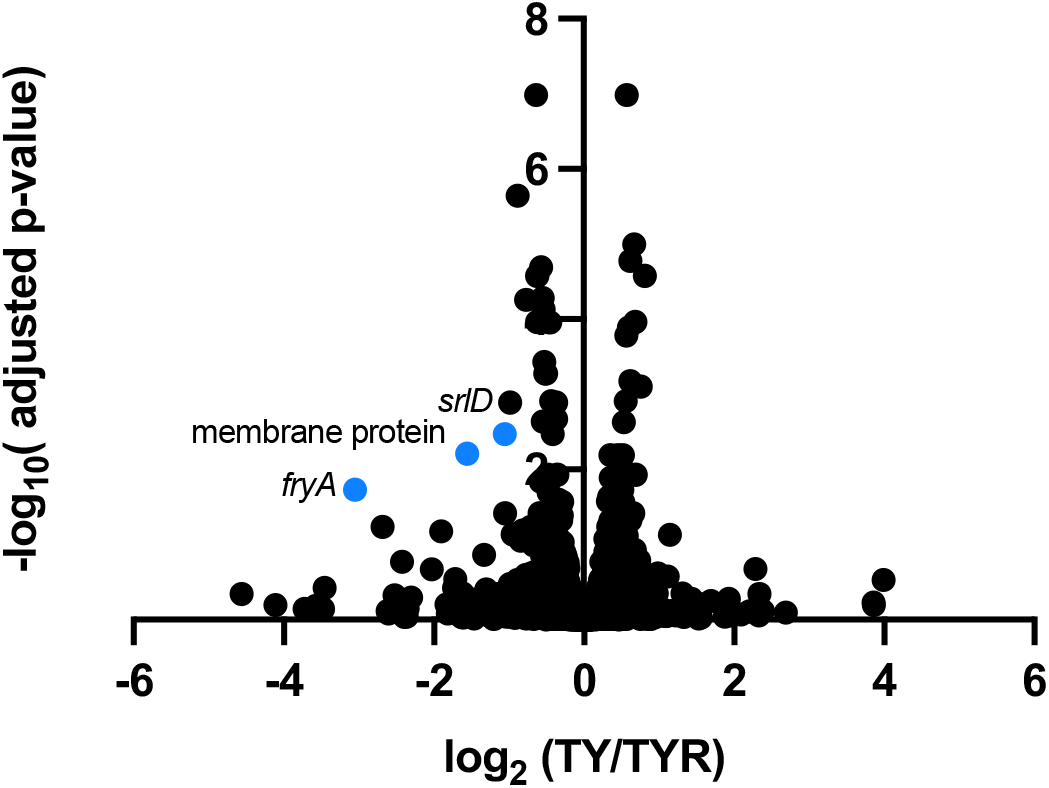
RNA sequencing of rhamnose-exposed *C. difficile*. Volcano plot of genes significantly altered by growth in tryptone-yeast extract(TY) media with 30 mM rhamnose in *C. difficile* VPI10643. Genes highlighted in blue have an adjusted p-value <0.05 and a log2 fold change greater than |1|. Adjusted p-values were generated using DEseq analysis. Negative fold-change indicates genes upregulated in TY-rhamnose(TYR) relative to expression in TY medium alone.

## Supplementary Tables

**Supplementary Table 1:** Patient metadata including EIA status and antibiotic exposure

**Supplementary Table 2:** Coefficients of logistic regression of antibiotic exposure associated with EIA staus

**Supplementary Table 3:** MaAsLin2 output of species associated with EIA status

**Supplementary Table 4:** MaAsLin2 output of metabolites associated with EIA status in addition to metabolite validation information

**Supplementary Table 5:** DEseq output of *in vitro C. difficile* transcriptomic profiling

